# Fluorescence imaging-induced phospholipid oxidation drives membrane phase separation by shifting miscibility boundaries

**DOI:** 10.64898/2026.09.11.750993

**Authors:** Angelina Duriez, Paige Tyra, Jacob Vehar, Alexander Olivero, Trevor Takubo, David A. Ford, Xiaohui Zha, Kevin C. Courtney

## Abstract

Fluorescence microscopy is a widely used tool for visualizing membrane organization in model and cellular systems; however, photoexcitation during imaging can generate reactive oxygen species, thereby altering membrane structure. Here, we show that fluorescence imaging actively drives membrane phase separation through photo-induced phospholipid oxidation. Using giant unilamellar vesicles, we observed that illumination of initially homogeneous membranes triggers the emergence of coexisting liquid-ordered and liquid-disordered domains. Chemical analysis by thin layer chromatography and mass spectrometry revealed the formation of a complex mixture of oxidized lipid species following excitation, characterized by stepwise oxygen additions to unsaturated phospholipids and multiple degradation products. These modifications increase lipid polarity and disrupt acyl chain packing, reducing favorable interactions with saturated phospholipids. As a result, membranes near the miscibility boundary become prone to demixing upon illumination, indicating that photo-induced lipid oxidation effectively shifts the miscibility boundary. Consistent with this model, light-induced phase separation is highly sensitive to membrane composition, with maximal effects observed near the liquid-liquid miscibility boundary. Electroformation substrate further modulated this effect, suggesting that pre-existing oxidative conditions sensitize membranes to subsequent photo-induced changes. Together, these findings demonstrate that excitation light can introduce artifacts in membrane phase behaviour. More broadly, subtle chemical modifications of phospholipids can shift membrane miscibility and reorganize membrane structure.

## Introduction

Biological lipid membranes are complex laterally heterogeneous bilayer structures where organization of proteins and lipids plays a central role in cellular functions, such as signaling transduction, and membrane trafficking(1-3). A key feature of lipid bilayer membranes, containing saturated and unsaturated phospholipids plus cholesterol, is the propensity to undergo liquid–liquid phase separation into coexisting ordered and disordered domains(4, 5). It is thought that proteins can be preferentially sequestered within these membrane domains, which facilitates protein sorting and promotes protein-protein interactions that are critical to cellular homeostasis(6-8). Lipid-mediated membrane organization has been extensively characterized using model membrane systems such as giant unilamellar vesicles (GUVs) and giant plasma membrane vesicles (GPMVs)(9-12). These model membranes provide a simplified experimental platform for dissecting the basic physicochemical principles underlying membrane phase behavior(11, 13). They have been essential in establishing how lipid composition controls miscibility, domain formation, and protein partitioning within membranes, enabling extrapolation of these organizational principles to complex biological membranes in live cells(14).

Fluorescence microscopy is one of the most widely used approaches for visualizing membrane organization in both model membranes and live cells. The incorporation of fluorescent lipid analogs into lipid bilayers enables direct observation of domain formation and dynamics with high spatial and temporal resolution(11). Several of these fluorescent lipids have been identified to preferentially associate with either ordered or disordered domains, making them a convenient tool for monitoring membrane phase separation in real-time(15). However, imaging by fluorescence microscopy is not an inherently passive process. For example, photoexcitation of fluorophores can generate reactive oxygen species, including singlet oxygen (^1^O_2_), through energy transfer to molecular oxygen(16-18). These reactive species can modify phospholipid structure, particularly those containing unsaturated acyl chains, leading to the formation of oxidized lipid derivatives with altered physicochemical properties(19, 20). Lipid peroxidation is known to influence membrane structure by increasing lipid polarity and perturbing acyl chain packing(19, 21-23). The addition of oxygen atoms on unsaturated hydrocarbon chains can further polarize co-existing saturated and unsaturated phospholipid mixtures, thereby impacting lipid miscibility(24, 25). It has been demonstrated that peroxide production during imaging can promote the formation of macroscopic membrane domains in GUVs (26-28). More broadly, lipid peroxidation was also recently shown to drive phase separation in GPMVs(24). Despite these reports, it remains unclear how widely appreciated this phenomenon is among researchers routinely using fluorescence microscopy to study membrane organization in both *in vitro* and live-cell contexts. Moreover, the extent of photo-induced lipid oxidation and its impacts on membrane properties remains incompletely understood. In particular, mechanistic characterization of oxidized lipid species and their links to membrane phase behavior remain limited. This raises an important concern, as unintended chemical modifications that are triggered by imaging could alter the very phenomena being studied.

In this study, we investigated the temporal impact of fluorescence illumination on membrane phase behavior in GUVs. We show that photon excitation can induce phase separation from homogeneous membranes and this effect is driven by the formation of oxidized lipid species. Using a combination of thin layer chromatography, mass spectrometry, and fluorescence imaging, we characterized the chemical nature of these oxidation products and link their formation to changes in membrane organization. Furthermore, we show that the extent of light-induced phase separation depends on membrane composition and proximity to the miscibility boundary, as well as experimental variables such as the choice of electroformation substrate.

Together, these findings elucidate photon excitation-induced lipid oxidation as a driver of membrane phase separation and emphasize an important source of experimental artifact in studies of membrane organization. These results provide insight into how chemical modifications of phospholipids can modulate membrane phase behavior and underscore the need for caution when using fluorescence imaging in both model membranes and live cells.

## Results

### Fluorescent imaging triggers membrane phase separation artifacts

While imaging GUVs composed of SM/DOPC/cholesterol (40:40:20) labeled with rhodamine-PE by fluorescence microscopy, we noticed that many of the initially homogenously labeled GUVs underwent phase separation during image acquisition (Fig. 1). Changes in temperature are a well-established driver of membrane phase separation(4, 29); however, all imaging in this study was performed at constant room temperature, highlighting a distinct phenomenon. This effect was also observed in complex protein-containing membranes derived from mammalian cells, i.e. GPMVs (Supplemental Fig. 1). Given the widespread use of GUVs and GPMVs to study membrane phase behaviour, these observations suggest that fluorescence imaging itself perturbs membrane organization, raising concerns about potential artifacts in microscopy studies of membrane phase separation.

**Figure 1.**
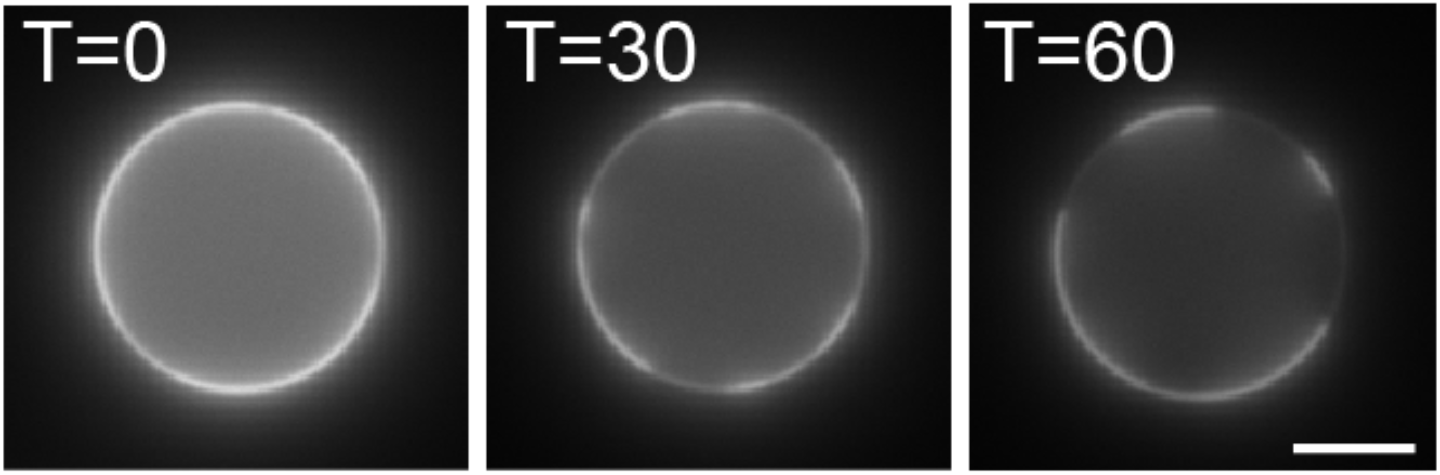
Representative images a GUV composed of DOPC/SM/cholesterol (40:40:20) labeled with 0.1% rhodamine-PE at the beginning (T=0), middle (T=30) and end (T=60) of continuous 60-second excitation regime at 560 nm. Scale bar represents 5 μm.

### Illumination generates oxidized lipid species

To better understand the principles driving light-induced phase separation, we sought to characterize the effects of fluorescence excitation. It has previously been reported that upon photoexcitation, fluorophores can form a longer-lived excited state that transfers energy to molecular oxygen (O_2_), generating singlet oxygen (^1^O_2_), a highly reactive oxygen species(16, 30). This ^1^O_2_ can then react with phospholipid acyl chains to create lipid hydroperoxides. We hypothesized that the addition of an oxygen group on the phospholipid would likely influence the packing of acyl chains within the membrane and provide a means to drive phase separation in membranes that would otherwise be miscible.

To confirm whether fluorescence excitation could indeed alter phospholipid structure, thus creating new lipid species, we first performed thin layer chromatography (TLC) on rhodamine-labeled SM/DOPC large unilamellar vesicles (LUVs) that were exposed to excitation light (560nm) in a fluorometer for 10 minutes. The lipids were then isolated from the resulting LUVs using the Bligh-Dyer extraction method and resolved by TLC(31). This analysis found that two new lipid species appeared following excitation, in comparison with the non-illuminated sample (Fig. 2). To identify the precursor of these new species, we repeated the TLC experiment using SM/Rho-PE and DOPC/Rho-PE LUVs separately, and confirmed that they originated from DOPC rather than SM (Supplemental Fig. 2). Next, to determine the nature of the two new lipid species that appeared on the TLC plate, we scraped the new bands from the TLC plate followed by lipid extraction and purification. These isolated lipids were then characterized by electrospray ionization mass spectrometry (MS) in negative ion mode by direct infusion of the samples in chloroform/methanol (1:4, v/v)(32). Four samples were analyzed: an unmodified DOPC control, DOPC exposed to 560nm light (prior to TLC separation), and the two unknown species isolated from the TLC plate following fluorescence excitation. The MS analysis found that the DOPC control sample contained a dominant peak of 820.55 *m/z*, indicative of a chloride adduct [M + Cl]^-^ (Fig. 3, top left panel). The additional molecular ions at 821.57 and 822.79 *m/z* in the control spectrum reflect the expected contribution of naturally occurring ^13^C isotopes to the 820.55 *m/z* ion. In agreement with our TLC experiment, we found the appearance of two new prominent molecular ions in the MS spectra of the illuminated DOPC sample, corresponding to 836.60 and 852.59 *m/z* (Fig. 3, bottom left panel). MS analysis of the isolated TLC bands confirmed that Band 1 (top) is dominated by the 836 *m/z* species, with smaller peaks at 837, 852, and 853 *m/z* and a dramatically reduced 820 *m/z* population (Fig. 3, top right panel), while Band 2 (bottom) is dominated by the 852 *m/z* species (Fig. 3, bottom right panel). Notably, these dominant 836 and 852 *m/z* species are 16 and 32 Da heavier than the unmodified DOPC, representing the incorporation of one and two oxygen atoms, respectively.

**Figure 2.**
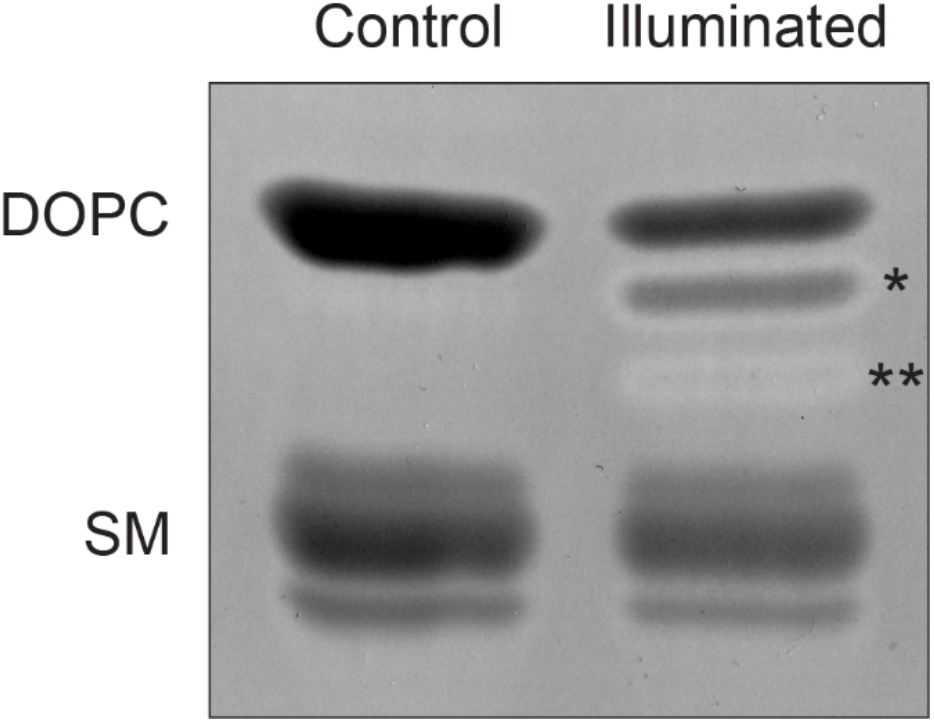
Representative thin layer chromatography of a DOPC and SM (1:1) lipid sample with 0.1% rhodamine-PE before and after exposure to 560 nm illumination in a fluorometer cuvette for 10 minutes. The single and double asterisk annotations indicate the appearance of two new bands that resolve in the sample following illumination.

**Figure 3.**
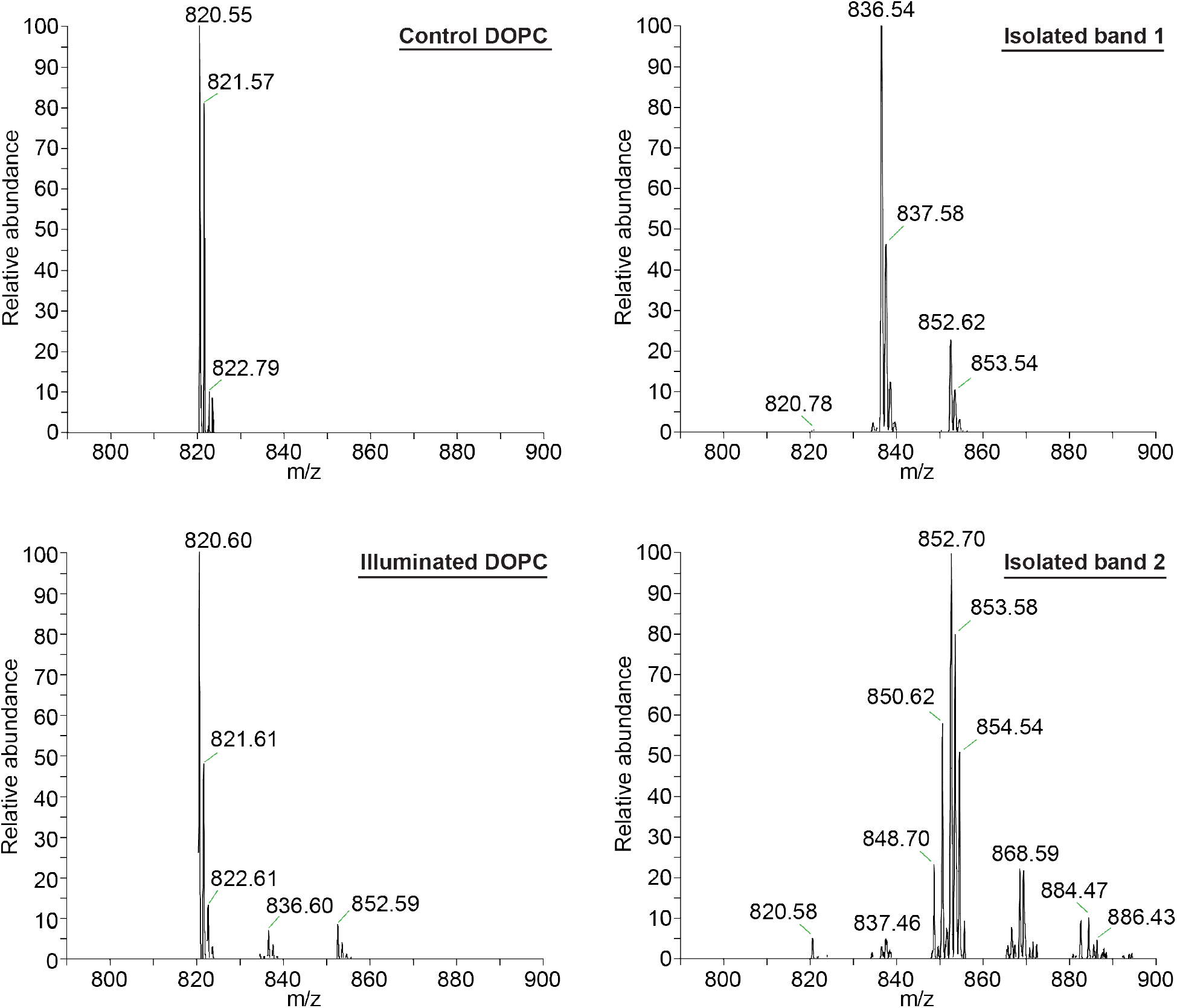
Mass spectrometry analysis of a DOPC sample with 0.1% rhodamine-PE before (Control DOPC) and after (Illuminated DOPC) illumination at 560 nm for 10 minutes in a fluorometer cuvette. The right panels represent lipids that were exposed to the light treatment, subsequently resolved on a TLC plate, as outlined in Figure 2, and isolated from the plate with a razor blade. Isolated Bands 1 and 2 correspond to the top and bottom bands from Figure 2, respectively.

Examining these spectra in more detail, we expect that the 836 *m/z* species corresponds to a mono-hydroxylated DOPC, in which one acyl chain bears a hydroxyl group (-OH), though a less likely isobaric epoxide structure (one oxygen bridging two neighbouring carbons) cannot be excluded by mass alone. The 852 *m/z* species is consistent with either a mono-hydroperoxide (one acyl chain bearing a hydroperoxide group) or an isobaric di-hydroxy structure (each acyl chain bearing a hydroxyl group); a mixture of both species is also plausible. Furthermore, the mass spectrum for isolated Band 2 shows a series of higher-order oxidation products at 868.59 and 884.47 *m/z*, corresponding to the addition of three and four oxygen atoms, respectively. Two additional peaks, at 850.62 and 848.70 *m/z*, are consistent with ketone-containing species: 850.62 *m/z* corresponds to a keto-alcohol structure (one acyl chain bearing a ketone, the other a hydroxyl group; +2 O, −2 H relative to unmodified DOPC), while 848.70 *m/z* corresponds to a di-ketone structure (both acyl chains bearing a ketone; +2 O, −4 H). Together, the appearance of these species (summarized structures depicted in Supplemental Fig. 3) shows that illumination leads to a complex mixture of secondary oxidative products that impede their migration across the TLC plate due to their increased polarity.

### Electroformation material influences light-induced phase separation

During the early stages of this work, we noted an anecdotal trend that GUVs formed on indium tin oxide (ITO)-coated slides are more prone to illumination-induced phase separation than those formed on platinum. This is consistent with an earlier study that found titanium electrodes were preferable to ITO electrodes for preventing peroxide production(26). ITO is a common GUV electroformation substrate due to being relatively inexpensive and for its transparent nature, allowing for active monitoring of the GUVs as they form. However, ITO is not chemically inert and, under electroformation conditions, may facilitate oxidative processes that contribute to lipid peroxidation artifacts(33). To systematically quantify the influence of the electroformation material on illumination-induced phase separation, we generated the same SM/DOPC/cholesterol GUVs using commercially available slides coated with either ITO, platinum or gold using a simple custom-built setup and imaged for 60 seconds by epi-fluorescence microscopy. We then quantified the number of GUVs that were initially phase separated in the full field of view in frame 1 of the video compared to the final frame at 60 seconds to determine the extent of light-induced phase separation triggered by the imaging regime. Note that the GUVs were initially identified by DIC prior to fluorescence imaging; they were only exposed to 560 nm light during the 60-second video acquisitions. We found that the initial phase separation present prior to our imaging regime was largely unchanged between the three materials. However, after the 60-second illumination period, we found that GUVs formed on the ITO slides had significantly more phase separated vesicles than those formed on platinum and gold coated slides (Fig. 4A). This suggests that GUVs electroformed on the ITO slides may be partially oxidized prior to imaging, which then becomes compounded by the subsequent 60-second imaging protocol.

**Figure 4.**
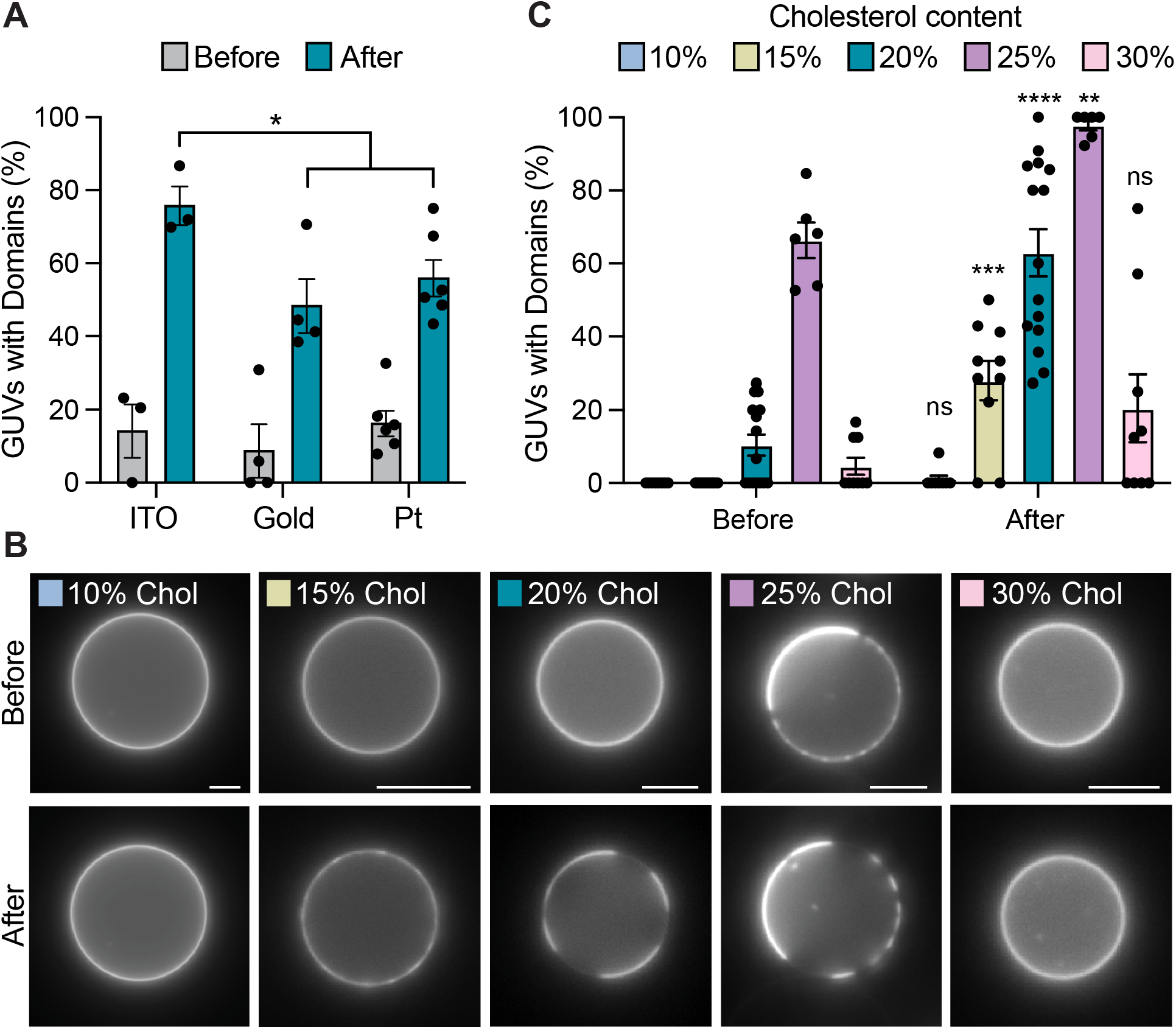
A) Quantification of phase separation in GUVs composed of DOPC/SM/cholesterol (40:40:20) with 0.1 % rhodamine-PE that were prepared on either ITO, gold or platinum coated glass slides. The number of GUVs exhibiting phase separation was measured before and after a 60-second imaging regime with 560 nm light exposure. GUVs prepared on ITO were compared to the pooled gold and platinum conditions by two-sided t-test; error bars represent standard error of the mean. B) Representative images of GUVs containing equimolar DOPC and SM with 10% - 30% cholesterol and 0.1% rhodamine-PE before and after exposure to 560 nm light for 60 seconds. Scale bar represents 10 μm. C) Quantification of GUV phase separation represented in panel B determined by one-way ANOVA with error bars representing standard error of the mean. The *, **, ***, and **** annotations represent P < 0.05, 0.005, 0.0005, and 0.0001, respectively.

### Illumination drives phase separation in lipid mixtures near the miscibility boundary

As described above, we initially hypothesized that the observed light-induced phase separation was a consequence of lipid peroxidation altering the acyl chain structure in the membranes to promote immiscibility. In this view, GUVs prone to light-induced phase separation would be at or near the immiscibility boundary for a ternary mixture and lipid peroxidation would serve to further polarize the DOPC from the SM in the presence of cholesterol to drive the phase separation. To validate this, we generated a series of lipid mixtures composed of equimolar DOPC and SM with increasing concentrations of cholesterol from 10-30%. We then generated GUVs using the platinum coated slides, again performed 60-second illumination acquisitions and quantified the extent of phase separation before and after illumination.

These experiments found that illumination markedly increased the fraction of GUVs exhibiting phase-separated domains in a cholesterol-dependent manner (Fig. 4B). Prior to the 60-second illumination regime, observable domain formation was minimal at low cholesterol concentrations (10-15%), modest at 20%, and most pronounced at 25%, with little domain formation observed at 30%. This suggests that 25% cholesterol is a composition at which intrinsic phase separation is most readily observed with equimolar DOPC and SM, as the majority of GUVs were phase separated from the outset without prior illumination with the 560nm laser. In contrast, the reduction in observable domains at 30% cholesterol likely reflects a membrane composition that favours a predominantly liquid-ordered phase, limiting the extent of visible ordered/disordered domain coexistence.

Following 60-second illumination, the prevalence of domain-containing vesicles increased across all compositions, with the largest effects observed at intermediate cholesterol levels (15-25%), which shifted from largely homogeneous membranes to predominantly phase-separated populations (Fig. 4B). The most dramatic response was observed at 20% cholesterol, where the phase-separated fraction rose from ∼10% to over 60% following illumination, while the 25% condition approached near-complete domain formation. These results indicate that photo-induced lipid oxidation shifts the miscibility boundary in a cholesterol-dependent manner, driving membranes near the phase transition into a phase-separated regime.

## Discussion

GUVs and GPMVs are widely employed model membrane systems used to elucidate principles of membrane phase separation(5, 13). These simplified membrane systems have been invaluable for describing the basic behavior of phospholipid bilayers and extrapolating phase separation phenomena in complex biological systems. It is well established that membranes containing saturated and unsaturated phospholipids can phase separate into ordered and disordered domains in a cholesterol-dependent manner(4). This principle is thought to promote myriad protein-protein interactions by enriching co-segregated proteins within ordered membrane domains; lipid-mediated protein organization has been demonstrated to drive directional cargo sorting in the secretory pathway and promote efficient signal transduction processes in the cell(2, 3, 6, 8).

However, in order to reliably apply the principles derived from model membrane systems to complex cellular contexts, stringent care must be taken to avoid experimental artifacts. The results presented here demonstrate that fluorescence imaging, a widely used method to characterize membrane properties, can actively perturb membrane organization by inducing phospholipid oxidation. This finding has important implications for the interpretation of fluorescence microscopy experiments using GUVs and GPMVs, as these techniques commonly rely upon fluorescent phase markers to monitor domain formation. It reveals that the act of observation itself can alter the physical state of the system under study.

Mechanistically, these data support a model in which fluorophore excitation generates ^1^O_2_, leading to oxidative modification of unsaturated lipid acyl chains. The mass spectrometry results clearly indicate the addition of oxygen atoms (+16, +32 Da, and higher-order products) after illumination. Given that ^1^O_2_ is known to add a hydroperoxide (-OOH) group on to acyl chain double bonds via the Schenck ene reaction, we expect the initial reaction to be a hydroperoxide addition, followed by the formation of a series of degradation products including alcohols (-OH) and ketones (=O) (Supplemental Figure 3)(34, 35). Hydroperoxides are inherently unstable and readily break down into these secondary products, rather than persisting as a stable end point(20). This degradation pathway accounts for the alcohol, keto-alcohol, and di-ketone species identified in the isolated bands, indicating that these arise as secondary products of an initially formed hydroperoxide rather than as independent primary oxidation events. The higher-order species observed at 868 and 884 *m/z* are consistent with a second, independent oxidation event on the opposing acyl chain, yielding molecules bearing oxidative modifications on both chains simultaneously. Moreover, the free radicals generated upon hydroperoxide decomposition can abstract hydrogen atoms from neighbouring lipid molecules, propagating a radical chain reaction that generates additional lipid radicals and new hydroperoxides(20). This chain propagation mechanism is expected to amplify oxidative damage beyond the initial ^1^O_2_-mediated reaction, contributing to the diversity of oxidized species observed.

Each of the chemical modifications that we observed are expected to disrupt the packing properties of DOPC by increasing polarity and reducing favorable hydrophobic interactions with neighboring lipids. In the context of ternary mixtures with sphingomyelin and cholesterol, such changes would reduce miscibility between oxidized DOPC and the more ordered SM-rich phase, thereby promoting separation. This provides a direct mechanistic explanation for how relatively small chemical modifications can produce large-scale changes in membrane organization.

Importantly, the compositional dependence of the oxidation effect reinforces this interpretation. The strongest light-induced phase separation was observed in mixtures near the miscibility boundary (20% cholesterol), where small perturbations in lipid interactions are expected to have the largest impact(29). In contrast, compositions far from the phase boundary showed more limited sensitivity to illumination. This behavior is consistent with a phase diagram shift rather than the creation of an entirely new phase regime, suggesting that lipid oxidation effectively shifts the thermodynamic landscape of the membrane. In practical terms, this means that fluorescence experiments probing phase behavior near critical points are particularly susceptible to photo-induced artifacts, requiring heightened experimental rigor. It should also be noted that photooxidation likely occurs to a similar extent across all membrane compositions, but its effects on lipid organization are only visually detectable near the miscibility phase boundary. Consequently, membranes far from the phase boundary may undergo equivalent photooxidation without any observable change in phase behavior, making illumination-induced damage especially pernicious in these cases, as it would go entirely undetected.

The choice of electroformation substrate also influences susceptibility to photo-induced phase separation. GUVs formed on ITO electrodes showed greater light-induced phase separation than those formed on platinum or gold, suggesting that ITO may introduce some degree of lipid oxidation during electroformation itself. This is consistent with Ayuyan and Cohen (2006), who found that GUVs electroformed on ITO electrodes contained measurable lipid peroxides, whereas GUVs formed on titanium electrodes were free of measurable peroxides(26). In this context, pre-existing oxidative damage would then prime the membrane for enhanced phase separation upon subsequent illumination. ITO, being relatively chemically reactive compared to noble metals, may facilitate lipid peroxidation during electroformation, lowering the threshold for phase separation triggered by illumination. This finding emphasizes that both imaging conditions and sample preparation can bias membrane behavior, potentially compounding experimental artifacts. In particular, this suggests that ITO-mediated electroformation may not be suitable for phase separation studies with GUVs. Moreover, the use of ITO-mediated electroformation could also have unintended consequences in proteo-GUV studies by altering protein function via oxidative damage.

Notably, lipid peroxidation proceeds via a self-propagating radical chain reaction, in which peroxyl radicals generated from an initial oxidation event abstract hydrogens from neighbouring lipids, generating new radicals and hydroperoxides(20). This propagating nature may help explain another trend we observed: while not systematically quantified in our study, GUVs imaged one day after electroformation showed substantially more phase-separated vesicles than those imaged the same day. Because oxidative damage can continue to propagate after its initial formation, pre-existing modifications from electroformation, ambient light exposure during storage, or potentially an earlier imaging session may compound over time, even without further active high-intensity illumination.

The observation that similar effects occur in GPMVs extends the relevance of these findings beyond simplified model systems. Lipid peroxidation was also recently reported by Balakrishnan and Kenworthy (2024) to have significant effects on phase separation in GPMVs(24). While GPMVs retain many aspects of native membrane complexity, they remain susceptible to photo-induced lipid oxidation and lack the antioxidant mechanisms of live cells, indicating that caution is warranted even in more biologically derived systems. This raises the possibility that oxidative artifacts could also influence interpretations of membrane organization in studies of intact cellular membranes, particularly when using high-intensity or prolonged fluorescence imaging of live cells.

These findings collectively underscore the need for careful experimental design in studies of membrane phase behavior. Minimizing illumination intensity and duration, selecting fluorophores with lower ROS generating potential, and validating results with non-fluorescent techniques are all important considerations(36).

Beyond their implications as an experimental artifact, the results of this work also suggest a broader conceptual link between lipid oxidation and membrane organization. In biological contexts, lipid peroxidation is a hallmark of oxidative stress and has been implicated in processes such as ferroptosis and neurodegeneration, as well as accumulation of oxidative damage during aging(37-39). The ability of lipid peroxidation to drive phase separation in model membranes raises the possibility that oxidative modifications could reorganize membrane domains *in vivo*, potentially affecting protein localization, signaling, and membrane trafficking. Such effects would be expected to perturb cellular homeostasis and contribute to disease development under increased oxidative stress. While the extent to which such mechanisms operate in cells remains to be determined, the present work provides a clear demonstration that oxidative chemistry can directly modulate membrane phase behavior.

In summary, this study identifies fluorescence-induced lipid oxidation as a driver of membrane phase separation and highlights an underappreciated source of experimental artifact in membrane biophysics studies. By linking photochemistry to shifts in membrane miscibility, these findings provide both a cautionary framework for interpreting imaging-based studies and a mechanistic basis for understanding how chemical modifications of lipids can regulate membrane organization.

## Methods

### Giant unilamellar vesicle electroformation

GUVs were created by electroformation using a simple setup consisting of a function generator and a BNC to dual alligator connector cable. 1 mM lipid mixtures of equimolar DOPC and sphingomyelin (mSM) plus 0.1% rhodamine-PE were prepared in chloroform with increasing cholesterol content ranging from 10-30 %. Lipids were deposited dropwise onto glass slides coated with either indium tin oxide, gold or platinum to form dotted lipid films across the surface, dried under a stream of nitrogen gas, and placed in a desiccator for at least one hour to remove residual organic solvent. A 250 μL volume of 200 mM sucrose solution was added to the lipid-coated region and contained using a greased O-ring. A second platinum slide was placed on top with conductive surfaces facing inward. The slides were connected to the function generator using alligator clips, and an alternating electric field (10 Hz, 3 V) was applied for 2 hours at room temperature in the dark. Following electroformation, the vesicle-containing solution was collected, filtered using a 3 μm membrane to remove impurities, and buffer exchanged into 25 mM HEPES, 100 mM KCl (pH 7.4).

### Fluorescence microscopy

Fluorescence imaging of GUVs was performed using a Nikon TE2000-E microscope in epi-fluorescence mode. The isolated vesicles were transferred and allowed to settle in an Ibidi bioinert 35 mm dish for a few minutes prior to imaging. Regions of interest were identified by low-light DIC imaging to avoid premature photoexcitation before the fluorescence excitation regime. For fluorescence imaging, GUVs containing rhodamine-PE dye were visualized with a 60x oil objective using fluorescence excitation at 550 nm and Cy3 emission filters. Fluorescence images were acquired once every second for 60 seconds under consistent illumination for all samples. Images before and after the imaging regime were analyzed using ImageJ software to assess membrane organization and domain formation.

### Thin layer chromatography

Lipids with and without light exposure were extracted from the aqueous buffer by the Bligh-Dyer method and loaded onto a 20x20 cm silica gel TLC plate using a 50 μL Hamilton syringe alongside purified lipid standards. The lipids were resolved on the plate by running in a chloroform/acetone/methanol/acetic acid/water (6:8:2:2:1) solvent system. The lipids were visualized by incubating the plate in a solution of 0.03 % Coomassie blue G, 30% methanol and 100 mM NaCl followed by destaining in 30% methanol and 100 mM NaCl.

To isolate the newly generated lipids species that were resolved on the TLC plate, the bands/silica were scrapped off the plate by a razor blade and collected on filter paper. The free silica was then transferred to a glass test tube and the lipids were extracted using the Bligh-Dyer method. The resulting isolated lipids were the characterized by electrospray ionization MS in negative ion mode as previously described(32).

---

**Supplemental Figure 1.**
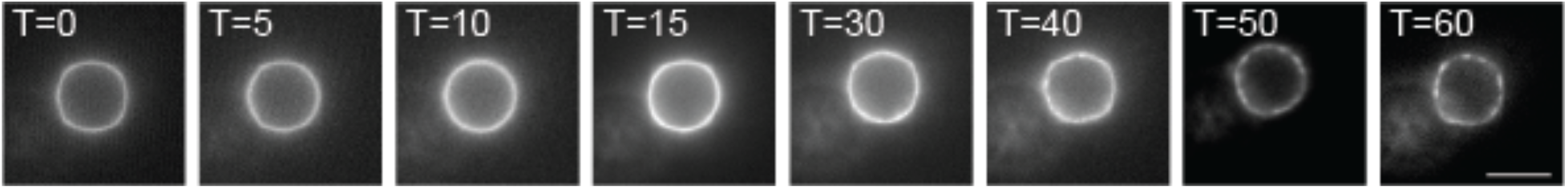
Representative images of a giant plasma membrane vesicle labeled with naphthopyrene that was exposed to 560 nm light for 60 seconds. Scale bar represents 5 μm.

**Supplemental Figure 2.**
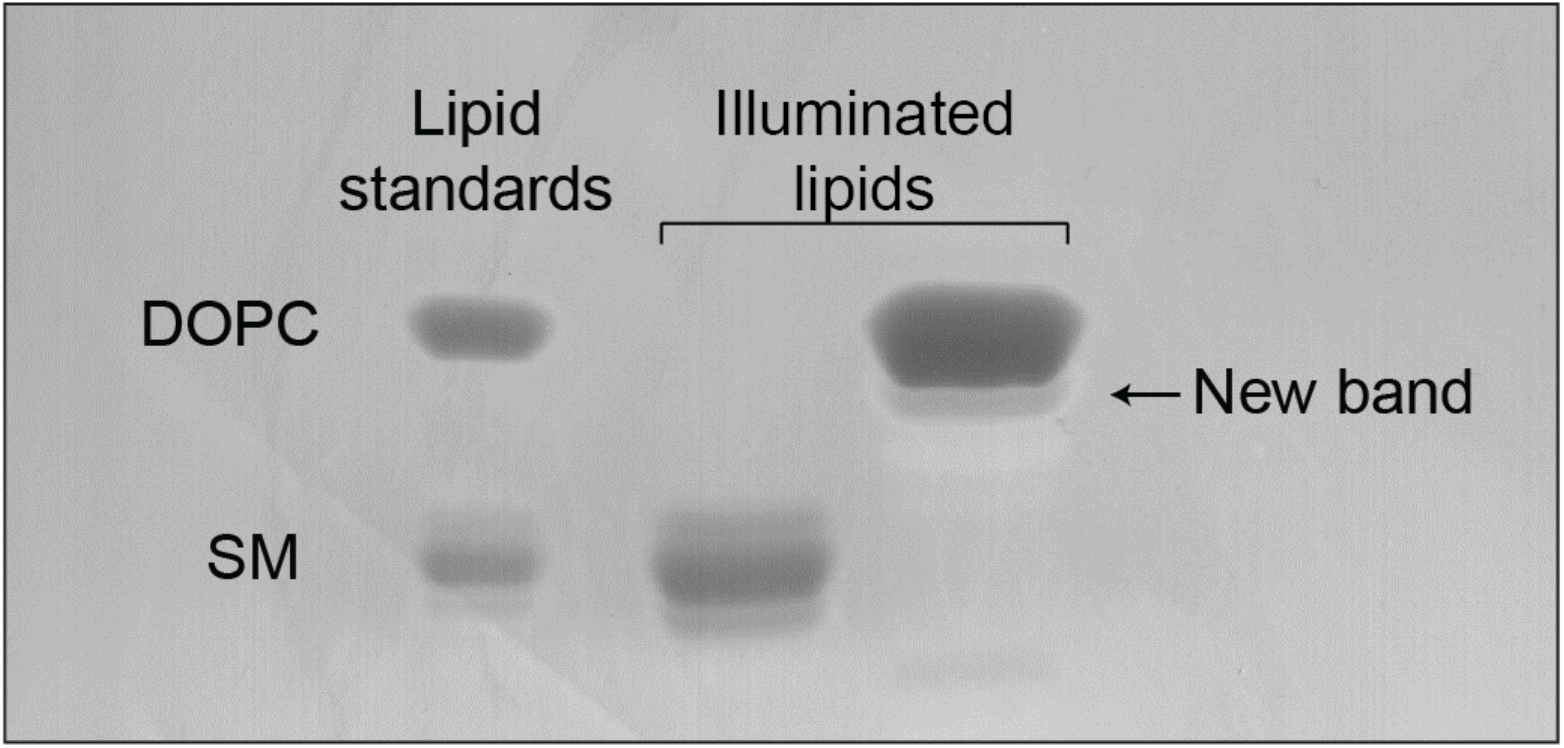
Thin layer chromatography of DOPC and SM with 0.1% rhodamine-PE exposed separately to 560 nm illumination in a fluorometer cuvette for 10 minutes.

**Supplemental Figure 3.**
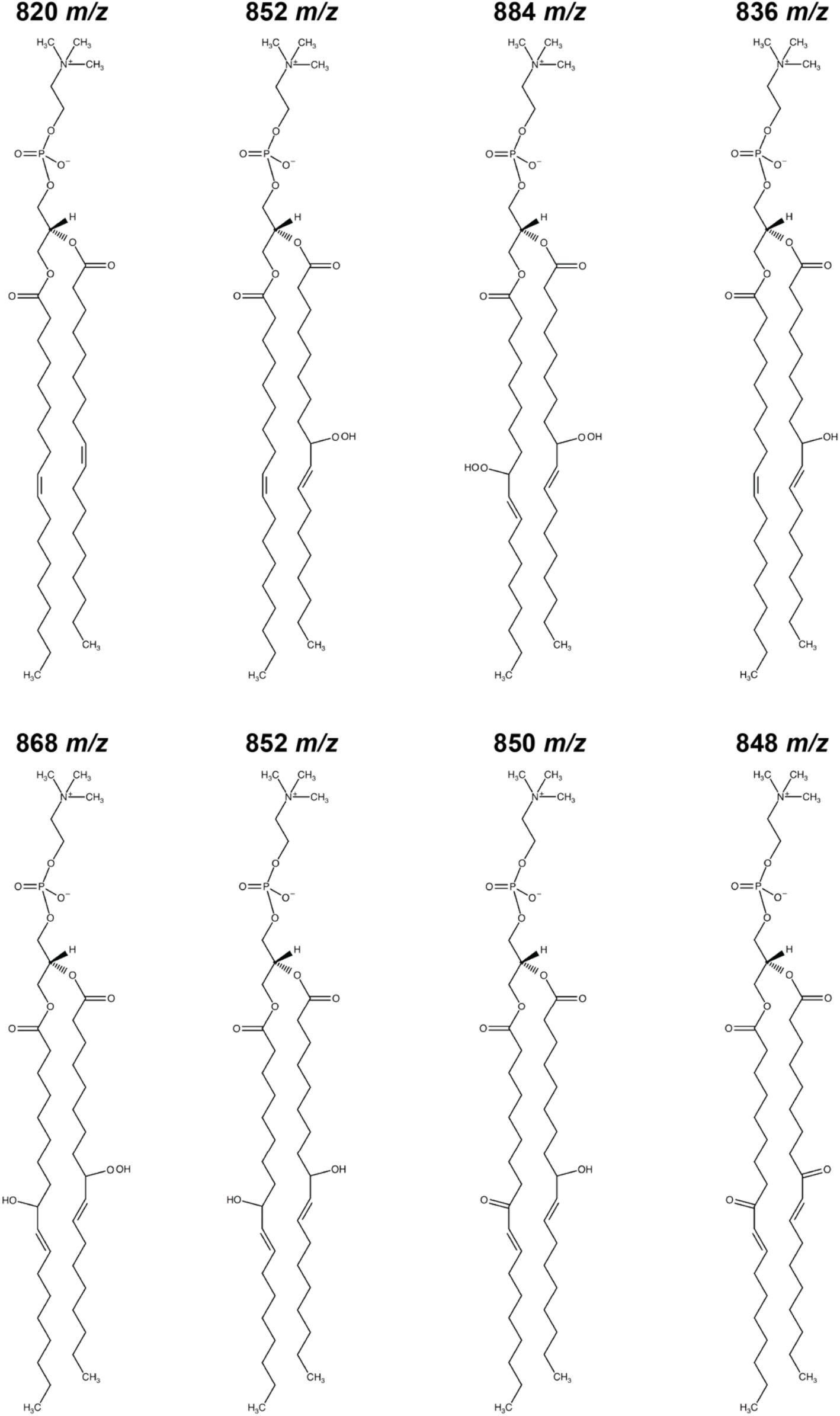
Summary of lipid species structures detected by mass spectrometry after illumination of a DOPC (820 *m/z*) sample containing rhodamine-PE with a 560 nm light source.

